# Relationship between thyroid hormones, their associated metabolites, and gene expression bioindicators in the serum of *Rana [Lithobates] catesbeiana* tadpoles and frogs during metamorphosis

**DOI:** 10.1101/2022.11.15.516608

**Authors:** Rikke Poulsen, Shireen H. Jackman, Martin Hansen, Caren C. Helbing

## Abstract

Anuran metamorphosis is characterized by profound morphological changes including remodeling of tissues and organs. This transition is initiated by thyroid hormones (THs). However, the current knowledge of changing levels of THs during metamorphosis relies on pooled samples using methods known for high variability with sparse reporting of measured variation. Moreover, establishing a clear linkage between key gene expression bioindicators and TH levels throughout the metamorphic process is needed. Using state-of-the-art ultra-high performance liquid chromatography isotope-dilution tandem mass spectrometry, we targeted 12 THs and metabolites in the serum of *Rana [Lithobates] catesbeiana* (n=5-10) across seven distinct postembryonic stages beginning with premetamorphic tadpoles (Gosner stage 31-33) and continuing through metamorphosis to a juvenile frog (Gosner stage 46). TH levels were related to TH-relevant gene transcripts (*thra*, *thrb*, and *thibz*) in back skin of the same individual animals. Significant increases from basal levels were observed for thyroxine (T4) and 3,3’,5-triiodothyronine (T3) at Gosner stage 41, reaching maximal levels at Gosner stage 44 (28±10 and 2.3±0.5 ng/mL, respectively), and decreasing to basal levels in juvenile frogs. In contrast, 3,5-diiodothyronine (T2) increased significantly at Gosner stage 40 and was maintained elevated until stage 44. While *thra* transcript levels remained constant and then decreased at the end of metamorphic climax, *thrb* and *thibz* were induced to maximal levels at Gosner stage 41, followed by a decrease to basal levels in the froglet. This exemplifies the exquisite timing of events during metamorphosis as classic early response genes are transcribed in anticipation of peak TH concentrations. The distinct T2 concentration profile suggests a biological role of this biomolecule in anuran postembryonic development and an additional aspect that may be a target of anthropogenic chemicals that can disrupt anuran metamorphosis and TH signalling. Hence, as a second aim of the study, we set out to find additional bioindicators of metamorphosis, which can aid future investigations of developmental disruption. Using a sensitive nanoLC-Orbitrap system an untargeted analysis workflow was applied. Among 6,062 endogenous metabolites, 421 showed metamorphosis-dependent concentration dynamics. These potential bioindicators included several carnitines, prostaglandins and some steroid hormones.

## 1 Introduction

Thyroid hormones (THs) are essential for the normal growth, development and health of all vertebrates (1–3) Among many other biological processes, THs control metabolism, cell differentiation and migration and also play a pivotal role for e.g., neurodevelopment (3,4). One of the most remarkable examples of TH action is anuran metamorphosis, where a tadpole transitions into a frog with extensive reprogramming and reorganization of organs and tissues (5–8). This remodeling has direct parallels to the developing human during the last trimester and the first weeks of life, where a surge of THs is essential for proper development (8–10). Hence, anuran metamorphosis is an excellent model system to study developmental endocrinology and the dynamics of the TH system. This has been acknowledged in several publications (5,7–9,11). The potential of amphibian metamorphosis as a screening tool for chemical disruption of the hypothalamus-pituitary-thyroid axis has furthermore been recognized in the OECD guideline 231 for the testing of chemicals i.e., the Amphibian Metamorphosis Assay (12).

Despite this, the current knowledge of baseline levels of THs during metamorphosis largely relies on pooled samples using methodologies known for high variability with sparse reporting of measured variation (13–23)(Table 2). In fact, (radio)immunoassays are still one of the most commonly used methods for quantification of THs despite known limitations, e.g., cross-reactivity and interference from proteins as summarized by Spencer (24). The current study uses a highly robust, selective and sensitive liquid chromatography tandem mass spectrometry (LC-MS/MS) method based on isotopic dilution to quantify 12 THs and metabolites (16,25).

THs exert their chemical messenger effect mainly through nuclear TH receptors, although growing evidence shows that non-genomic signaling also plays a role (8,26). In the genomic pathways, nuclear TH receptors (THRs) bind directly to DNA or indirectly by interacting with chromatin associated proteins to modulate expression levels of hormone-responsive genes (26). Two THR genes, a and b, code for these receptors. Both have been found to show specific expression patterns according to cell type and developmental stage (27). In addition to TH concentrations, the transcription levels of THR genes therefore serve as robust indicators of TH signaling and the endocrinological processes during metamorphosis. Earlier studies have shown that an abundance of transcripts encoding THR a and b (*thra*, *thrb*) as well as TH-induced bZip protein (*thibz*, Fig. 1F) show high correlation with developmental stage in anuran tadpoles (27–29) indicating their potential to serve as bioindicators of metamorphosis in addition to THs. However, these data and their relationships to changes in TH levels *in the same subjects* have not been established in amphibians.

**Figure 1.**
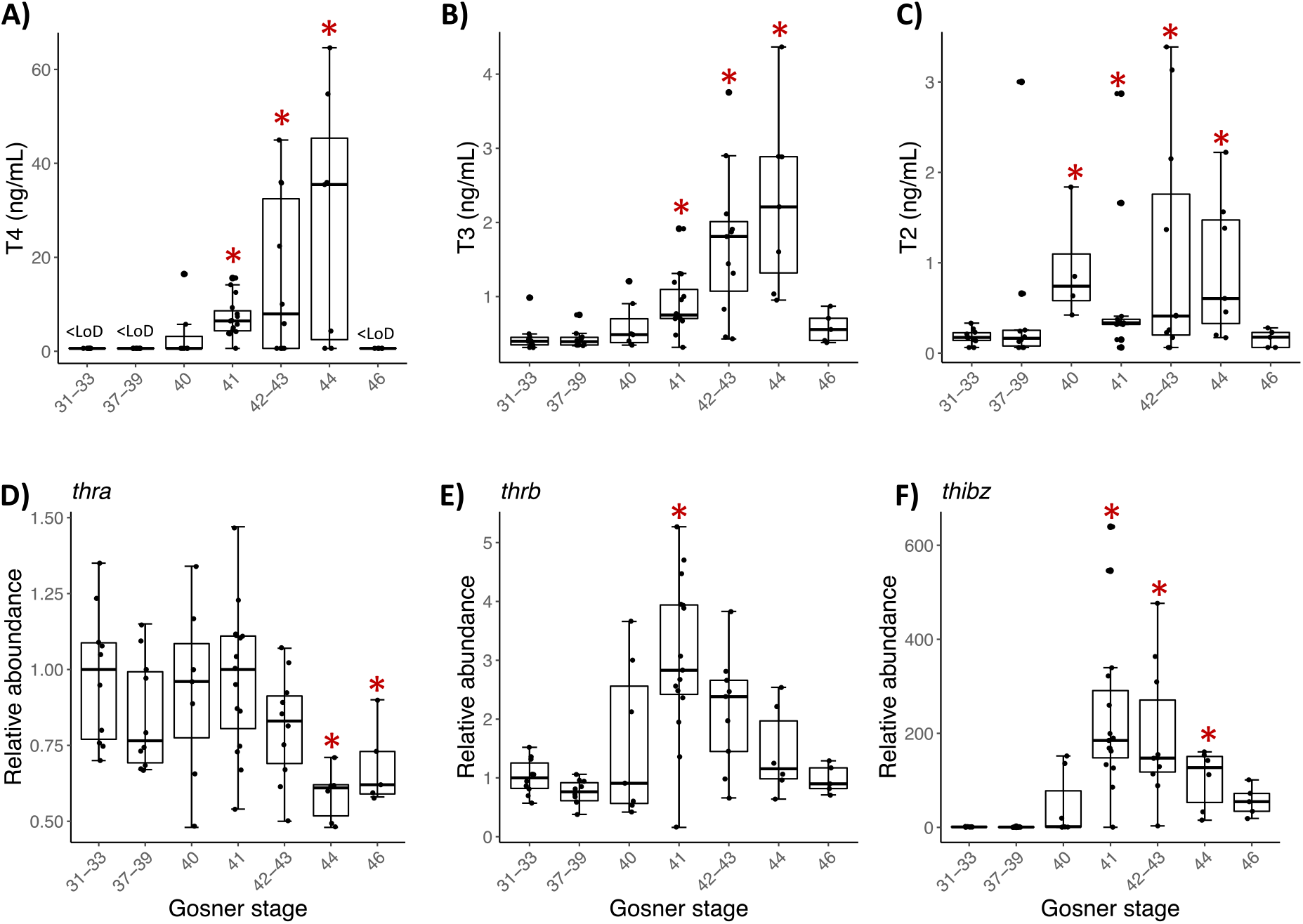
Box plots showing serum concentration (ng/mL) of (A) T4, (B) T3, and (C) T2 and back skin gene expression of (D) thra, (E) thrb, and (F) thibz for the grouped Gosner stages. The lower and upper hinges correspond to the first and third quartiles. The whiskers extend from the hinge to the largest/smallest value no further than 1.5 times the inter-quartile range. Circles show individual replicates. Asterisks indicate significant difference from the premetamorphic Gosner stage 31-33 (Dunn’s post hoc test, p-value <0.05).

American bullfrog (*Rana [Lithobates] catesbeiana*) is a great alternative model species as it is a representative of the largest amphibian group, Ranids or “true frogs”, and is found worldwide. Their large-sized tadpoles enable the analysis of individual animals rather than pooled samples. Furthermore, this amphibian compared to e.g., *Xenopus laevis* tadpoles, also has a greater similarity to human life stages as they completely transition from aquatic to terrestrial living (*Xenopus* remain fully aquatic as adults). By using wild caught animals instead of laboratory manipulated individuals, we ensure that the natural variation of endocrinological state is represented. By using a high sample replication we also capture the natural biological variation without losing statistical power.

Although anuran metamorphosis is initiated solely by THs, many other biochemical pathways undergo extensive remodelling during this transition (30) and the regulation of metamorphosis is a complex process of interacting mechanisms (11). The complexity, tight regulation, and interconnection of biological networks and pathways can be conceptualized using untargeted metabolomics. In contrast to traditional hypothesis-driven investigations, untargeted approaches, such as metabolomics, interrogate the whole biological system and let the organism-response guide the investigations (31–35). Such knowledge is very powerful when trying to make new discoveries about a biological system and can aid the search for new robust biological markers. In the case of anuran metamorphosis, this type of analysis provides a holistic view of the biological processes and allows for a search of robust bioindicators of the molecular dynamics during the transitions. Such bioindicators are useful for assessing the state of the endocrine system and its disruption by external environmental factors (such as endocrine disrupting chemicals or climate change (i.e., temperature)).

The first aim of the present study is to map the dynamics of known bioindicators, ie. TH concentration and transcription of TH sensitive genes in wild-caught *R. catesbeiana* tadpoles and frogs during metamorphosis. This includes the quantification of measurable THs and metabolites in serum as well as abundance of gene transcripts of *thra, thrb*, and *thibz* in back skin of the same animals. As a second aim, we use untargeted metabolomics to reveal new bioindicators of metamorphosis.

## 2 Materials and methods

### 2.1 Sampling

*Rana [Lithobates] catesbieana* tadpoles of unknown sex were caught locally in Victoria, British Columbia, Canada and housed in the University of Victoria Outdoor Aquatics Unit. Animals were maintained with a 12 h light/12 h dark photoperiod in 340 L fiberglass tanks containing 15°C recirculated and dechlorinated municipal water and fed daily with Spirulina flakes (Dynamic Aqua Supply, Surrey, BC, Canada). Treatment of animals was in accordance with the guidelines of the Canadian Council on Animal Care and experimental protocol #2015-028 was approved by the University of Victoria Animal Care Committee.

Tadpoles and recently metamorphosed froglets were euthanized at distinct postembryonic developmental stages (36). The mean ± standard error weights and number of animals are shown in Table S1(36). Animals were euthanized in 0.1% (w/v) tricaine methanesulfonate (Syndel Laboratories Ltd., Vancouver, BC, Canada) buffered with 25 mM sodium bicarbonate in dechlorinated municipal water. Blood was collected by tail muscle incision for tadpoles and severing of the aorta for froglets. The collected blood was allowed to coagulate at room temperature for 10-15 min. The blood from each animal was kept separate and never pooled. The coagulated blood was centrifuged at 10,000 *xg* for 10 minutes at 4°C, the serum was collected, immediately flash frozen in liquid nitrogen and stored at −80°C until use. On average, 150 μL serum per individual animal was collected using this procedure.

The back skin from the same animals was harvested from each individual tadpole or frog from between the eyes to the tip of the rostrum to permit the most consistent comparison of skin tissue across developmental stages. The skin was immediately immersed in RNAlater (Invitrogen, Thermo Fisher Scientific) and stored at −20°C until RNA extractions were performed.

### 2.2 Gene expression bioindicators

RNA extraction, cDNA synthesis, and quantitative real time polymerase chain reaction (qPCR) analyses were performed as previously described (37). Three normalizer gene transcripts encoding ribosomal protein L8 (*rpl8*), ribosomal protein S10 (*rps10*), and elongation factor 1 alpha (*eef1a*) were used, and three classical TH-responsive transcripts were analysed encoding thyroid hormone receptor α (*thra*), thyroid hormone receptor β (*thrb*), and thyroid hormone induced basic region leucine-zipper containing transcription factor (*thibz*). The exact run conditions and primer sequences have been previously reported (29,37,38).

### 2.3 TH and metabolite extraction

Serum samples were extracted for targeted TH determination as previously described (39) and untargeted metabolomics using the same extract. In brief, 50 μL serum was spiked with isotopic-labelled (^13^C)-thyroid hormone standards (internal standards (IS), cT2, cT3 and cT4) and homogenized. After antioxidant treatment (100 μL, 25 mg/mL ascorbic acid, R,R-dithiothreitol and citric acid solution) and protein denaturation by urea (8 M in 1% ammonium hydroxide), the samples were enriched using solid-phase micro-extraction (SOLAμ HRP 10 mg/1 mL 96 well plate, ThermoFisher Scientific, Germany) and reconstituted in 100 μL 5% methanol containing an instrument control standard (ICS, crT3). Procedural blanks containing water instead of plasma were included from the beginning. Detailed descriptions of chemicals and reagents are found in the supplementary material (S1.1).

### 2.4 TH quantification

The targeted analysis of THs was performed on an Agilent 6495c triple-quadrupole system with a hyphenated Agilent 1290 Infinity II ultra-high performance liquid chromatography (UHPLC) system (binary pump, degasser, and autosampler; Agilent Technologies) as previously described (39). Targeted analytes were thyroxine (T4), 3,3’,5-triiodothyronine (T3), 3,3’,5’-triiodothyronine (rT3), 3,5-diiodothyronine (T2), 3,3’-diiodothyronine (3,3’-T2), 3-iodothyronine (T1), thyronine (T0), 3-iodothyronamine (T1Am), 3-iodothyroacetic acid (T1Ac), 3,5-Diiodothyroacetic acid (Diac), triiodothyroacetic acid (Triac) and tetraiodothyroacetic acid (Tetrac). LoDs as fmol on column for all THs were 4.0 (T4), 0.79 (T3), 0.76 (rT3), 0.56 (T2), 0.60 (3,3’-T2), 0.74 (T1), 0.23 (T0), 0.05 (T1Am), 9.4 (T1Ac), 15 (Diac), 10 (Triac), 9.6 (Tetrac). Neat standard ten-point equimolar calibration curves (0.04 – 20.0 pmol/mL TH, n=2) were prepared in 5 % methanol and all vials contained a fixed amount IS and ICS (15.2 pmol/mL). Data analysis was conducted in MassHunter version 10.1 (Agilent Technologies). Detectable hormones in our samples were T4, T3, T2, T1Am and T0. Statistical analysis and visualization of the gene expression biomarkers and TH concentrations were done using R ver. 3.6.3 (40). The univariate statistical analysis was performed by Kruskal-Wallis test followed by pairwise comparison by Dunn’s test (41). To allow for statistical comparisons, TH levels below the limit of detection (LoD) were replaced by ½ LoD. Data between the LoD and limit of quantification (LoQ) remained as they were. The data was visualized using ggplot2 (42). Principal component analyses (PCA) were done using prcomp and visualized by ggbiplot (43).

### 2.5 Untargeted metabolomics

The untargeted analysis was performed on a nanoflow UHPLC Orbitrap mass spectrometer system (ThermoFisher Scientific) with a preconcentration trap-column setup described elsewhere (33). The instrument was operated in data-dependent mode by automatically switching between MS and MS/MS fragmentation. Samples were analyzed in randomized order and a pooled sample was injected in between every five samples to correct for any instrumentational fluctuations and monitor any systematic errors. Quality and validity of the analysis was confirmed by principal component analysis (PCA) showing that the composite quality control (QC) samples were centrally located in the plot (Fig. S2). Compound Discoverer (CD) software version 3.3 (Thermo Scientific) was used for data processing and analysis as previously described (33).

A univariate analysis (one-way ANOVA with Tukey as post-hoc test and p-values adjusted by Benjamin-Hochberg algorithm) was performed in Compound Discoverer (CD) to filter out compounds of particular interest to differentiate between developmental stages. Using R ver. 3.6.3, abundance of selected compounds was visualized in a heatmap using the R-package pheatmap (44) with Euclidian clustering on rows and no clustering on columns. A multivariate analysis by sparse partial least squares discriminant analysis (sPLS-DA) was performed on the 421 selected compounds using the R-package mixOmics (45) in R ver. 4.3.1 (40). Putatively annotated compounds were based on comparisons of the experimental MS2-spectrum with in-house spectral libraries (annotation level 1) and the online spectral library, mzCloud, (annotation level 2)(46). Putatively characterized compound classes (annotation level 3) were based on assigned predicted composition, *in silico* fragmentation by SIRIUS4 (47), and plausible, tentative candidates to that composition. For instance, whether the compound was likely to naturally occur in tadpole serum. Compounds that did not meet these criteria were annotated to level 4 and not incorporated in the functional analysis. The functional analysis was based on the Kyoto Encyclopedia of Genes and Genomes (KEGG) database(48), BioCyc(49), and human metabolome database (HMDB)(50).

## 3 Results

### 3.1 Thyroid hormone profile and gene expression bioindicators

Of the targeted THs T4, T3, T2, T0 and T1Am were above the instrumental limit of detection. Metamorphosis-related concentration dynamics was observed for T4 (Fig. 1A), T3 (Fig. 1B) and T2 (Fig. 1C). The concentration of T0 and T1Am did not change significantly over the recorded developmental window (Figure S1), except for a transient decrease in T0 around metamorphic climax in stage 41-44 compared to premetamorphic stage 31-33 (Fig. S1A).

The metamorphosis related concentration pattern was similar for T4 and T3. Until Gosner stage 40, T4 was below the limit of detection (LoD, 1.3 ng/mL), after which the concentration gradually increased reaching a maximum of 28 ± 10 ng/mL at Gosner stage 44 (Table S2) resulting in a 20-fold increase. In the froglets (Gosner 46), T4 was again below the detection limit. T3 was quantifiable in all samples and, like T4, increased gradually during metamorphosis, reaching the highest concentration at Gosner stage 44 (2.3 ± 0.5 ng/mL) before decreasing to premetamorphic levels in froglets. The difference between premetamorphosis and metamorphic climax was about 5-fold. T2 concentrations were generally lower with higher variation (Figure 1A). However, in contrast to T3 and T4, the concentration of T2 was already significantly higher at stage 40 compared to the initial pre-metamorphic measurement in Gosner stage 31-33 animals (Dunn’s post hoc test p-value =0.002). This higher level was maintained at about 5-fold through metamorphic climax at Gosner stage 44 before returning to basal levels in the froglet.

Back skin gene expression of the thyroid hormone receptor β (*thrb*, Fig. 1E) and TH-induced bZip protein-encoding transcripts (*thibz*, Fig. 1F) also showed the expected correlation with Gosner stage. Both transcripts increased quickly in abundance from Gosner stage 40, peaked at Gosner stage 41, and then gradually decreased to basal levels in the froglet (Gosner stage 46). The change was particularly noticeable for *thibz;* abundance increased around 500-fold compared to Gosner stage 40. In comparison, the change in *thrb* was only 3-fold. Thyroid hormone receptor a (*thra*, Fig. 1D) showed a stable abundance in these back skin samples until stage 44, where it decreased to a significantly lower level compared to the premetamorphic group (Gosner stage 31-33). The fold change was, however, minor.

As animal-specific TH quantification and gene expression was obtained, a multivariate analysis by PCA was performed to observe whether the developmental stages would cluster across the two principal components of the data (Fig. 2). Driven mainly by the concentration of T3 and T4 as well as the *thibz* and *thrb* transcript abundance there is a division along component 1 (55% explained variation) that separates the pre- and early prometamorphic stages (Gosner 31-40) plus the juveniles (Gosner 46) from the animals at the late prometamorphic stage and metamorphic climax (Gosner 41-44). The direction of the vectors represents the correlation with the principal component and the length represents the contribution to the component. The vector of T4 is nearly parallel to the X axis, indicating that the difference of PC1 is mainly due to the contribution of this TH. Furthermore, it is mainly *thra* abundance that drives the separation on principal component 2.

**Figure 2.**
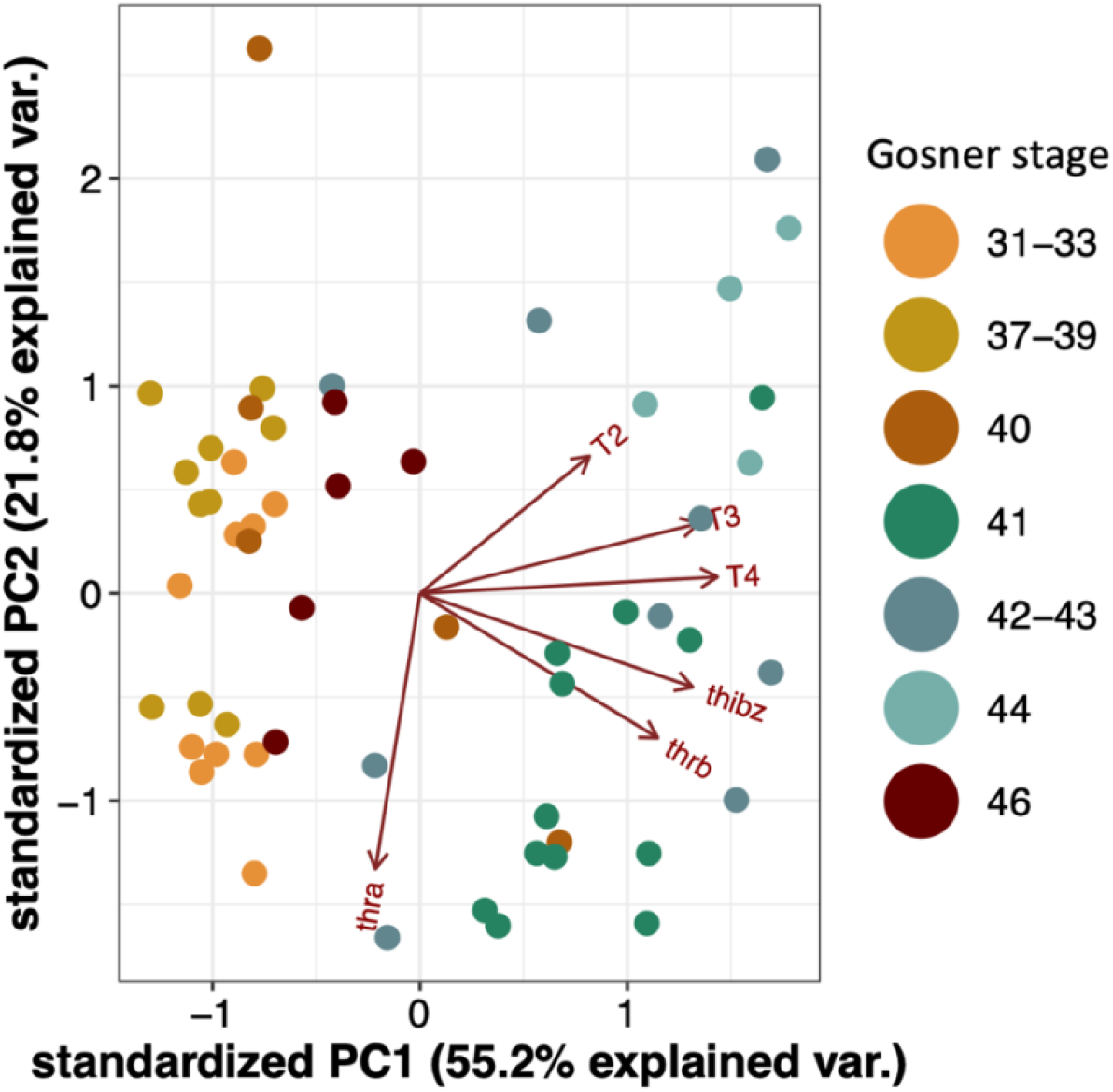
Principal component analysis (PCA) biplot for T4, T3, and T2 concentrations and gene transcript abundance of thra, thrb and thibz. Dots represents samples from individuals and colors represents Gosner stages.

We also investigated the individual Spearman’s rank correlations between gene transcript abundance and TH concentrations and found a strong and highly significant correlation for T4 and T3 versus *thibz* (r>0.6, p-value < 0.0001) and *thrb* (r>0.4, p-value < 0.001). T2 was also weakly correlated with *thrb* and *thibz* (r>0.3, p-value = 0.01). For all correlations see supplementary Table S3.

### 3.2 Untargeted metabolomics

After filtering out background peaks from more than 200,000 features, 6,062 endogenous metabolites remained. The measured metabolites were dominated by compounds active in lipid metabolism including fatty acids, bile acids, and steroid hormone derivatives. A good coverage of arachidonic acid metabolism and linoleic acid metabolism was observed.

Based on the observation that T4 and T3 reached a maximum concentration at Gosner stage 44, and that this was significantly different from the concentration in premetamorphosis and in juveniles, we wanted to look for metabolites that showed similar concentration dynamics. A differential analysis was performed comparing Gosner stage 44 to premetamorphic stages (Gosner 31-39) and juveniles (Gosner 46). This resulted in 421 significant metabolites (p-value_adj_<0.05). As exemplified by Figure 3A, most of them followed the curve shape of T4 (Fig. 1A) and T3 (Fig. 1B), including a gradual increase up to a maximum reached at metamorphic climax, after which the level dropped down again at the froglet stage. Examples of compounds are given to the right including arachidonic acid (Fig. 3B), a prostaglandin (Fig. 3C) and a hydroxyeicosatetraenoic acid (Fig. 3D), which are all related to the arachidonic acid metabolic pathway. The latter two, especially, show a pattern similar to the THs and exhibit high fold changes. In contrast to previous observations (30), arachidonic acid does not follow this trend in the data set. Rather, there is considerable variation in measurements with a substantial reduction in the juvenile (Fig. 3A). Dodecenoylcarnitine (Fig. 3E) and palmitoyl glutamic acid (Fig. 3F) are other examples of compounds that follow the pattern of the THs. Carnitines are involved in the transport of long chain fatty acids into mitochondria for beta-oxidation and energy generation. Palmitoyl glutamic acid is a palmitic acid amide of glutamic acid and belongs to the compound class known as N-acylamides.

**Figure 3.**
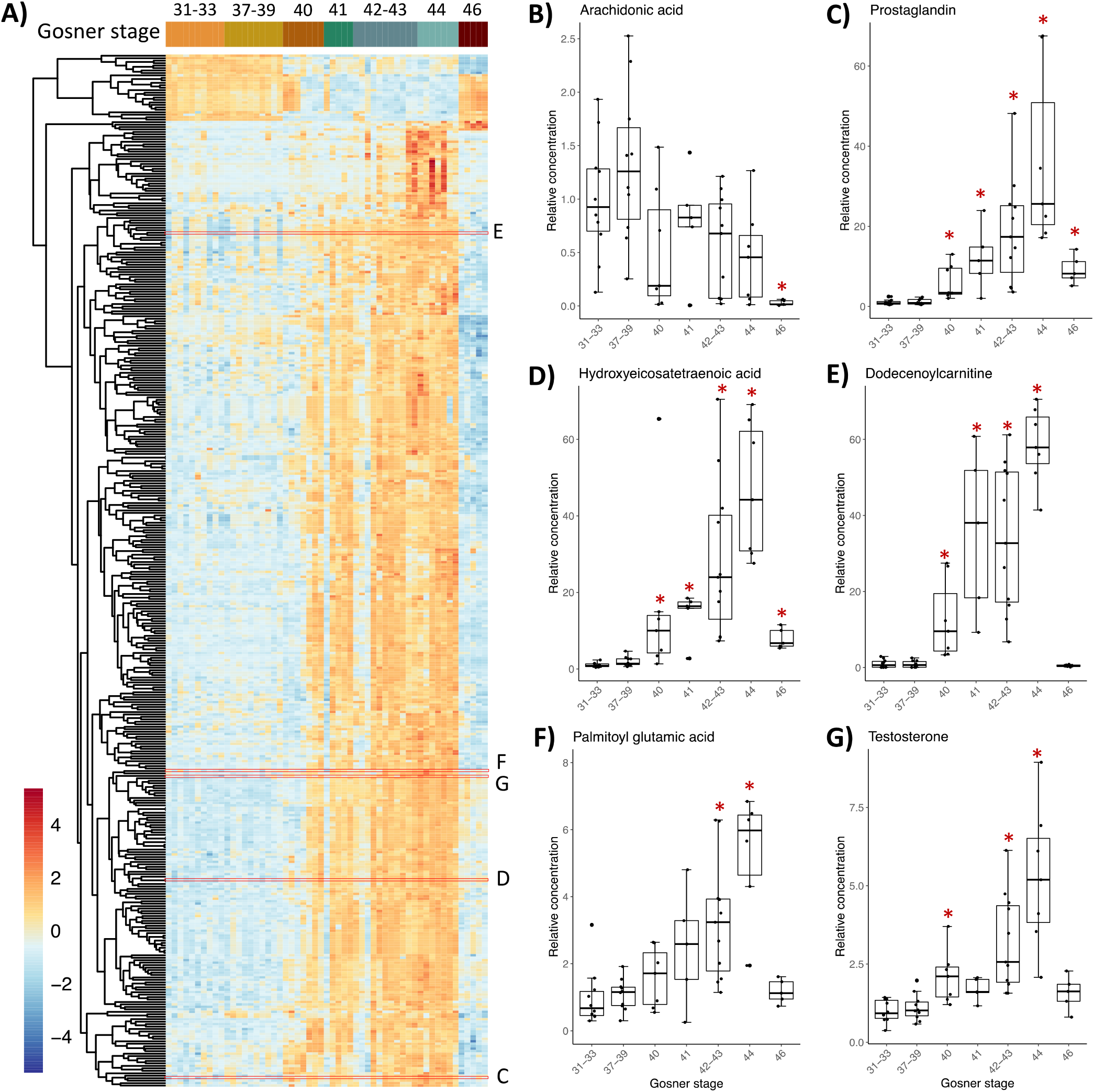
Heatmap (A) of 421 compounds that differed significantly in relative concentrations (ANOVA, adj p-value <0.05) between Gosner stage 44 and either stages 31-33 (premetamorphosis) or stage 46 (juveniles). Boxplots (B-G) showing examples of metabolites with their relative concentration plotted against the grouped Gosner stages. The positions of metabolites appearing in the heatmap with corresponding boxplots are outlined by red boxes. The relative concentration is found by normalizing peak areas to the average peak area for Gosner 31-33. Asterisks indicate significant difference from the premetamorphic Gosner stage 31-33 (Dunn’s post hoc test, p-value <0.05).

All 421 compounds may be efficient bioindicators of metamorphosis and possibly for its disruption. An sPLS-DA analysis was performed to discover compounds that best discriminate between Gosner stages (Figure S3A). The first component explained 45% of the variation and metabolites, which contributed to this first component, mainly separated the prometamorphosis and metamorphic climax, where T4, T3, and T2 levels were maximal. After tuning the model, 20 variables (i.e., metabolites) were selected on component 1 and all the selected variables had the highest abundance in Gosner stage 44 animals (Fig. S3B). The compounds included dodecenoylcarnitine (Fig 3E) and decanoylcarnitine both identified with a spectral match (level 1). Table 1 shows 12 of these compounds, for which a chemical class could be assigned, including their stability (i.e., the proportion of cross validation folds (across repeats) where it was selected to be used for component 1). The details for all 20 compounds can be found in Table S4.

**Table 1.** Annotated metabolites (n=12) which contributed to the first component in the sPLS-DA analysis. Chemical class, ID level (46) and name (tentative for all ID-level<1) is given as well as their sPLS-DA stability score (i.e., the proportion of cross validation folds across repeats where it was selected to be used for component 1). Further details are provided in Table S4.

| Chemical class | ID level | Name | sPLS-DA Stability score | Formula |
| --- | --- | --- | --- | --- |
| Acyl carnitines | 1 | Dodecenoylcarnitine | 0.83 | C19 H35 N O4 |
| Acyl carnitines | 3 | Undecylcarnitine | 0.67 | C18 H35 N O4 |
| Acyl carnitines | 3 | (5E)-tetradecenoyl-L-carnitine | 0.57 | C21 H39 N O4 |
| Acyl carnitines | 1 | Decanoyl-L-carnitine | 0.4 | C17 H33 N O4 |
| Prostaglandin | 3 | Prostaglandin C1 | 0.37 | C20 H32 O4 |
| Linoleic acids and derivatives – Fatty acyls | 3 | 9-OxoOTrE | 1 | C18 H28 O3 |
| Linoleic acids and derivatives – Fatty acyls | 3 | 2R-HpOTrE | 0.43 | C18 H30 O5 |
| Linoleic acids and derivatives | 3 |  | 0.5 | C18 H28 O4 |
| Long-chain fatty acid | 3 |  | 1 | C18 H28 O3 |
| Long-chain fatty acid | 3 |  | 0.77 | C18 H28 O3 |
| Medium-chain hydroxy acids and derivatives | 3 |  | 0.47 | C19 H32 O5 |
| Medium-chain hydroxy acids and derivatives | 3 |  | 0.87 | C19 H32 O5 |

## 4 Discussion

The present study brings forth essential new knowledge of baseline levels of THs and TH-responsive genes during metamorphosis. Currently, published data largely rely on pooled samples, limited replication, and sparse reporting of variation (Table 2). In contrast, the present study includes individual sampling of 5-10 animals per developmental stage and the methodology reflects state-of-the-art quantification of biomolecules. This also includes the quantification of TH metabolites other than T3 and T4, such as T2, which provides novel insight into the thyroid hormone system. Table 2 summarizes maximal levels of THs (ng/mL) measured during anuran metamorphosis in published studies. The average reported value for all species is 6.7±0.6 ng/mL and 2.0±0.6 ng/mL for T4 and T3, respectively. The studies of *R. catesbeiana* show slightly lower average values of 6.0±0.9 ng/mL and 1.1±0.9 ng/mL for T4 and T3, respectively. In contrast, the concentrations measured in the present study are higher. As shown in the final row of Table 2, we report maximum levels of 28±10 ng/mL and 2.3±0.5 ng/mL for T4 and T3, respectively. The reason for this dissimilarity could reflect differences the developmental stage that was sampled for quantification (see Gosner stage in Table 2) and the masking of variability by using pooled samples. Additionally, the extraction method can cause differences especially for T4. T4 is largely protein-bound (51), and to obtain a quantification of total T4, the extraction method in the present study included a protein denaturation by urea to release all protein-bound hormone. This differs from earlier applied methodologies and could explain the higher quantity measured in the present study. Notably in the only two previous studies examining T4 levels in individual tadpoles, it was noted that all stages had some individuals where T4 was not detected (20,21). In the present study, we found Gosner stage 44 animals all had detectable T4 concentrations but two individuals had T4 levels below the LoQ. The present study results are, therefore, consistent with these earlier findings in that T4 is highly variable, which contributes to the high variation in T4 levels during metamorphic climax.

**Table 2.** Summary of maximal levels of THs (ng/mL) measured during metamorphosis in published studies. All previous studies used pooled samples with two exceptions where measured samples were from individuals (20, 21). The final row lists the values measured in the present study. RIA, radioimmunoassay; SEM, standard error of the mean; UHPLC, ultrahigh-performance liquid chromatography; UPLC, ultra-performance liquid chromatography; ID, isotope dilution, MS/MS tandem mass spectrometry

| Species | Gosner Stage | Tissue | Assay Used | T<sub>4</sub> (ng/mL)<br>Mean±SEM | T<sub>3</sub> (ng/mL)<br>Mean±SEM | T<sub>2</sub> (ng/mL)<br>Mean±SEM | Reference |
| --- | --- | --- | --- | --- | --- | --- | --- |
| *R. catesbeiana*<sup>a</sup> | 45 | Plasma | RIA | 5.0±0.3 | 0.8±0.1 | - | (18) |
| *R. catesbeiana*<sup>a</sup> | 45 | Plasma | RIA | 4.6±0.5 | 0.8±0.2 | - | (18) |
| *R. catesbeiana* | 43 | Plasma | RIA | 5.0±0.4 | 1.4±0.1 | - | (13) |
| *R. catesbeiana* | 43-44 | Plasma<sup>b</sup> | RIA | 6.0 | 1.5±0.3 | - | (19) |
| *R. catesbeiana* | 43-45 | Serum | RIA | 9.4±0.6 | - | - | (20) |
| *R. clamitans* | 45 | Plasma | RIA | 7.2±0.5 | - | - | (21) |
| *R. esculenta* | 44 | Serum | RIA | 2.6 | 0.1 | - | (22) |
| *X. laevis* | 44 | Plasma | RIA | 7.5 | 5.2 | - | (23) |
| *X. laevis* | 44 | Serum | RIA | 5.4 | 0.2 | - | (22) |
| *X. laevis* | 44 | Plasma<sup>c</sup> | UPLC-ID-MS/MS | 10.7±0.8 | - | - | (14) |
| *X. borealis* | 42-44 | Plasma | RIA | 8.7 | 4.5 | - | (15) |
| *X. clivii* | 42-44 | Plasma | RIA | 9.2 | 2.0 | - | (15) |
| *X. muelleri* | 42-44 | Plasma | RIA | 6.4 | 4.0 | - | (15) |
| *R. catesbeiana* | 44 | Serum | UHPLC-MS/MS | 28±10 | 2.3±0.5 | 0.9±0.3 | The present study |

<sup>a</sup>Two data sets were provided in Regard et al, 1978
<sup>b</sup>From ethanol extract
<sup>c</sup>From a single sample

In addition to updated information on circulating thyroid hormones and metabolites (T4, T3, T2, T0 and T1Am), this is also the first study to simultaneously analyse same-individual back skin gene expression of the thyroid hormone receptors α and β (*thra*, *thrb)* and TH-induced bZip protein (*thibz*) throughout metamorphosis. Efforts have been made earlier to combine published data on *X. laevis* to get this simultaneous view (8,52), but the same individual measurement, with high level of replication offers an unprecedented depiction of the endocrinological dynamics during anuran postembryonic development, and by approximation, vertebrate development as well. In previously published illustrations, the peak in T3 concentration was reported to occur simultaneously with the peak in *thrb* transcript abundance and precede the maximum in T4 (23,52). In the present study, however, we observed that the peak in T3 and T4 occurs simultaneously, and that *thrb* abundance preceded the increased concentration of both T3 and T4. This exemplifies the exquisite timing of events during the metamorphic process as the synthesis of the receptor occurs in anticipation of the peak in TH concentration.

The observed timewise correlation between THs and the abundance of *thibz* and *thrb* was confirmed by correlation analyses. Furthermore, the PCA showed that T3 and T4 as well as *thrb* and *thibz* transcript abundance were the main factors to separate the metamorphic climax from other developmental stages. Because the back skin transcript data are so well linked to the TH measurements it opens up the possibility of using non-lethal skin swabs on the rostrum area as a validated proxy for TH levels to detect the status and the possible disruption of the TH system.

The abundance of *thra* transcripts did not, however, follow the same pattern; its abundance remained stable until the end of metamorphic climax where it decreased significantly and remained low in the juveniles. This significant reduction coincides with maximal T4/T3 and it could potentially be an example of negative feedback. The observed metamorphosis-related dynamics is indeed in good correspondence with an earlier study in *X. laevis*, where a plateau in *thra* expression also was observed until a drop after metamorphic climax (53). Relative THR expression is, however, known to be tissue specific, so it is important to note that our results illustrate the transcript abundance in back skin, a tissue that remodels but is retained throughout metamorphosis. Other tissues may exhibit different dynamics as exemplified by the differences in tissue responsiveness to THs (29) and tissue specific expression profiles of THRs (54). In particular, *thra* seems to have a very diverse expression pattern across tissues (27,29,54).

To the best of our knowledge, this is the first study to quantify T2 throughout tadpole metamorphosis. Interestingly, T2 increased earlier than T3 and T4 prior to the significant increase in transcript abundance of *thrb* and *thibz* and remained high during metamorphosis. This indicates a specific and distinct biological role of this biomolecule in anuran postembryonic development. There is limited research on the action of T2 in different vertebrates and, to the best of our knowledge, there are no studies about the biological role in amphibians. In humans and rodents it has, however, been established that this iodinated molecule has biological activity distinct from that of T4 and T3 (55), including for instance activation of fatty acid oxidation and down-regulation of lipogenesis in hepatic fatty acid metabolism (56–61). In the freshwater fish, tilapia, T2 acts as a selective activator of a specific long isoform of THR-b named L-TRb1 and modulates its expression *in vivo* (58). T2 also specifically regulates gene networks associated with cell signaling and transcriptional pathways in brain and liver, which are distinct from those affected by T3 (59,60). Furthermore, T2 affects myelination in cerebellum (59), which is essential for neurodevelopment and participates in growth processes via L-TRb1 activation (61).

Based on these findings and the fact that T2 concentration was significantly increased earlier than T3 and T4, it is tempting to speculate that T2 may modulate THR expression in amphibians. More research is clearly needed to investigate this hypothesis. Generally, TH-related research tends to focus on T4 and T3, of which the biological role is well-documented. However, deiodinated products like T1 and T2, as well as aminated metabolites (e.g., T1Am) and iodothyroacetic acids (e.g., 5-diiodothyroacetic acid, Diac), are increasingly being recognized as biologically active (57,62,63). These molecules will be an important focus in furthering our understanding of TH dynamics.

The untargeted analysis showed that the changes in THs during metamorphosis are accompanied by similar dynamics in many other biomolecules. Earlier untargeted studies in the serum of *R. catesbeiana* using ultraperformance liquid chromatography coupled with mass spectrometry (UPLC-MS) have found a high degree of remodelling in five core metabolic pathways during metamorphosis. This included arginine and purine/pyrimidine, cysteine/methionine, sphingolipid, and eicosanoid metabolism and the urea cycle (30). In addition, a major role for lipids during the postembryonic process was established and arachidonic acid metabolism also underwent remodeling during metamorphosis (30). In accordance with this, the current study also observed metamorphosis-related changes in several lipids including eicosanoids and arachidonic acid metabolites (Figure 3). The bioindicators that showed the highest predictive power for distinguishing metabolite dynamics like the THs, however, included prostaglandins, acyl-carnitines and fatty acyls (Table 1).

L-Carnitine is an endogenous metabolite found in all mammalian species (64). The most important role of L-carnitine is to facilitate the transport of fatty acids into the mitochondria for beta-oxidation (64). During this process, L-carnitine forms a high-energy bond with a fatty acid to give an acyl carnitine derivative that is transported into the mitochondrial matrix. Here it undergoes transesterification with coenzyme A (CoA) to form the corresponding acyl-CoA, which then enters the fatty acid b-oxidation (64). In this way, L-carnitine is also able to regulate the intracellular ratio of free CoA to acyl-CoA by forming acylcarnitines and eventually remove surplus of acyl groups from the organism by preferential renal excretion. This means that the intramitochondrial relationship between free CoA and acyl-CoA is reflected in the extracellular L-carnitine to acylcarnitine ratio, giving indirect evidence of altered mitochondrial metabolism (64). Considering this, our observation of altered carnitine levels during metamorphosis, which additionally correlates with the TH-dynamics, is consistent with the fact that the mitochondrial metabolism is modified during metamorphosis. A tissue specific metabolic reorganization during metamorphosis has already been observed for *R. omeimontis* tadpoles (65). This study found that, by metamorphic climax, these tadpoles showed overall increased energy metabolism. In the liver, this was accompanied by upregulated amino acid catabolism and downregulated β-oxidation and glycolysis, suggesting a metabolic shift from carbohydrate and lipids to amino acid. A similar shift is likely occurring in *R. catesbeiana*. Therefore, the results in the current study show that the acyl-carnitines are excellent bioindicators for the metabolic shift during metamorphosis.

The ratio of L-carnitine to acyl-carnitines is currently applied in neonatal screening for inborn errors of metabolism, so the potency of these bioindicators is already known from other vertebrates (66). Acyl carnitine concentrations and THs can be biochemically linked as T3 is a known inducer of carnitine palmitoyltransferase I gene expression in the liver (67). This enzyme catalyzes the transfer of long chain fatty acyl groups from CoA to carnitine for translocation across the mitochondrial inner membrane. A long chain fatty acyl, which was tentatively annotated as 2R-HpOTrE, and its oxidation product, 9-OxoOTrE, were actually also among the 20 compounds selected on sPLS-DA component 1. The effect of THs on the transferases for short and -medium chained fatty acyls to the best of our knowledge, not been investigated. However, the two metabolites that were classified as medium-chain hydroxy acids and derivatives (Table 1) could potentially be related to this.

We also observed modifications to the arachidonic acid metabolism pathway and eicosanoids during metamorphosis (Figure 3). Furthermore, a prostaglandin was among the 20 compounds that best discriminate between Gosner stages (Figure 3C, Table 1). Prostaglandins are synthesized by sequential enzymatic modification of arachidonic acid (68). They are mainly recognized for their role in inflammatory responses (69) but they also play a role in neurogenesis (70). A link to TH homeostasis have been observed in rats with high iodide intake-induced hypothyroidism where increased levels of prostaglandins were found (71). Therefore, prostaglandins and other eicosanoids present an additional chemical class of important bioindicators of metamorphosis and vertebrate developmental endocrinology that could be applied in future studies.

The steroid hormone, testosterone, was interestingly found to follow a similar Gosner stage profile as T3, T4, carnitines and prostaglandins. Testosterone is the main sex hormone in vertebrate males. However, because the animals in the study were of mixed sex, testosterone seems to serve a different purpose in this developmental window. It should be noted that the observed increase in testosterone occurred after anticipated sexual differentiation and is completely independent of sexual maturation. Much empirical evidence exists that confirms crosstalk between the hypothalamic-pituitary-thyroid (HPT) axis which involves TH synthesis and the hypothalamic-pituitary-gonadal axis which involves synthesis of steroid hormone (7,72). As reviewed in Thambirajah et al. (7) this includes the effect of androgens like testosterone on the HPT axis. The exact mechanisms and physiological consequences are still unclear; however, androgens have been linked to bone formation and growth (73), which may be the reason for the observed metabolic pattern during metamorphosis.

The vital crosstalk between the HPT axis and the hypothalamic-pituitary-adrenal axis, which signals the production of corticosteroids, is also well established. This includes hormones like cortisol and corticosterone. Corticosterone is maximal at metamorphic climax much like T3 and T4 and it can accelerate TH-dependent metamorphosis. It has also recently been shown to also be required for survival through metamorphosis in *X. tropicalis* (74). A metabolite tentatively annotated as corticosterone was found among the compounds that displayed same Gosner stage profile as T3 and T4, however it could not be annotated with a spectral match. These findings underline what many others has called for before, namely that crosstalk between endocrine axes receives more attention in future studies. It is one more piece in the puzzle that is essential for our understanding of vertebrate developmental endocrinology. The non-target data provides a necessary context for interpreting TH-mediated developmental processes and potential molecular crosstalk mechanisms. Furthermore, this provides a solid foundation from which alternative bioindicators of metamorphosis may be chosen as feasible endpoints in future risk assessment of chemicals and their potency for causing developmental toxicity.

## 6 Author Contributions

**Rikke Poulsen**: Conceptualization, methodology, investigation and formal analysis, writing – original draft. **Shireen H. Jackman**: Methodology, investigation and formal analysis, writing – review and editing. **Martin Hansen**: Conceptualization, methodology, writing – review and editing, funding acquisition and resources. **Caren C. Helbing**: Conceptualization, methodology, investigation and formal analysis, writing – review and editing, funding acquisition and resources.

## 8 Acknowledgments

We thank Sara Ohora and Emil Egede Frøkjær for technical assistance and Dr. Nancy Denslow for helpful discussions

## 9 Data Availability Statement

The metabolomics dataset generated for this study can be found in the MassIVE database under the accession number MSV000090714

