## supplementary material for "Relationship between thyroid hormones, their associated metabolites, and gene expression bioindicators in the serum of *Rana [Lithobates] catesbeiana* tadpoles and frogs during metamorphosis"


### 1 Animal data

**Table S1. Mean, standard error (SEM), and number of animals used in the present study.** Serum and back skin were collected from each individual for analysis.

| Gosner stage | Number of animals | Weight±SEM (g) |
|---|---|---|
| 31-33 | 10 | 13.91±0.76 |
| 37-39 | 10 | 16.57±1.17 |
| 40 | 7 | 20.47±4.60 |
| 41 | 15 | 29.07±1.68 |
| 42-43 | 6 | 25.70±2.72 |
| 44 | 14 | 24.54±1.62 |
| 46 | 5 | 14.45±1.32 |

### 2 Thyroid Hormone Analysis (Expanded)

### 2.1 Chemicals, reagents and materials

Thyroxine (T4, CAS 51-48-9), 3,3',5-triiodothyronine (T3, CAS 6893-02-3 ) and 3,3',5'-triiodothyronine (rT3, CAS 5817-39-0) as well as ($^{13}$C)-thyroid hormone standards $^{13}C_6$-L-thyroxine (cT4, CAS 720710-30-5), $^{13}C_6$-3,3',5-triiodothyronine (cT3, CAS 1213431-76-5) and $^{13}C_6$-3,3',5'-Triiodo-L-thyronine (crT3, CAS 1786403-77-7) were obtained from from Supelco® through Merck (Darmstadt, Germany) as solutions (100 µg/mL in methanol with 0.1N NH3, Cerilliant®) with purity ≥99% and isotopic purity ≥98.0%, respectively. 3,5-diiodothyronine (T2, CAS 1041-01-6, purity ≥99%), 3,3-diiodothyronine (3,3'-T2, CAS 4604-41-5, purity 98%), thyronine (T0, CAS 1596-67-4, purity ≥ 98.0 %), triiodothyroacetic acid (Triac, CAS 51-24-1, purity ≥90%), tetraiodothyroacetic acid (Tetrac, CAS 67-30-1, ≥ 98.0 %), 3-iodothyronamine (T1Am, CAS 788824-64-6, purity ≥ 98.0 %) and 13C6-3,3´-diiodo-L-thyronine (cT2, CAS 1217459-13-6, purity 99 atom % 13C, 97%) were obtained from Sigma-Aldrich® through Merck (Darmstadt, Germany). 3-iodothyronine (T1, CAS 10468-90-3, purity 98%), 3-iodothyroacetic acid (T1Ac, CAS 60578-17-8, purity 98%), 3,5-Diiodothyroacetic acid (Diac, CAS 1155-40-4, purity 98%) were obtained from Santa Cruz Biotechnology (Dallas, Texas, USA). Stock solutions of solid compounds were prepared in methanol (Optima® LC-MS grade, Fisher Scientific) and diluted in high-purity water (Optima® LC-MS grade, Fisher Scientific).

The reagents L-ascorbic acid (CAS 50-81-7, purity ≥99%), R,R-dithiothreitol (CAS 3483-12-3, purity 97%), citric acid (CAS 77-92-9, purity 99%) and urea (CAS 57-13-6), purity 99.5% ) were obtained from Sigma-Aldrich, Germany. All solvents used for extraction and analysis were of Optima® LC-MS grade obtained from Fisher Scientific

### 2.2 Thyroid hormone results

**Table S2. Concentration (ng/mL) of quantified THs (T4, T3, T2, T0 and T1Am) as mean ± standard error of the mean (SEM) for each group of Gosner stages.**

| Gosner stage | T4 (ng/mL) Mean ± SEM | T3 (ng/mL) Mean ± SEM | T2 (ng/mL) Mean ± SEM | T0 (ng/mL) Mean ± SEM | T1Am (ng/mL) Mean ± SEM |
|---|---|---|---|---|---|
| 31-33 | <LoD* | 0.45 ± 0.06 | 0.18 ± 0.03 | 0.1415 ± 0.0004 | 0.047 ± 0.001 |
| 37-39 | <LoD* | 0.43 ± 0.04 | 0.48 ± 0.29 | 0.1420 ± 0.0007 | 0.045 ± 0.001 |
| 40 | 3.6 ± 2.3 | 0.60 ± 0.12 | 0.94 ± 0.31 | 0.142 ± 0.002 | 0.048 ± 0.001 |
| 41 | 7.2 ± 1.1 | 0.90 ± 0.10 | 0.60 ± 0.22 | 0.145 ± 0.007 | 0.047 ± 0.003 |
| 42-43 | 15.8 ± 5.5 | 1.71 ± 0.30 | 1.06 ± 0.38 | 0.152 ± 0.012 | 0.051 ± 0.005 |
| 44 | 28.1 ± 10.0 | 2.28 ± 0.46 | 0.94 ± 0.30 | 0.1384 ± 0.0004 | 0.044 ± 0.001 |
| 46 | <LoD* | 0.58 ± 0.09 | 0.16 ± 0.04 | 0.194 ± 0.035 | 0.066 ± 0.013 |

**LoD = Limit of detection. For T4 LoD = 1.3 ng/mL*

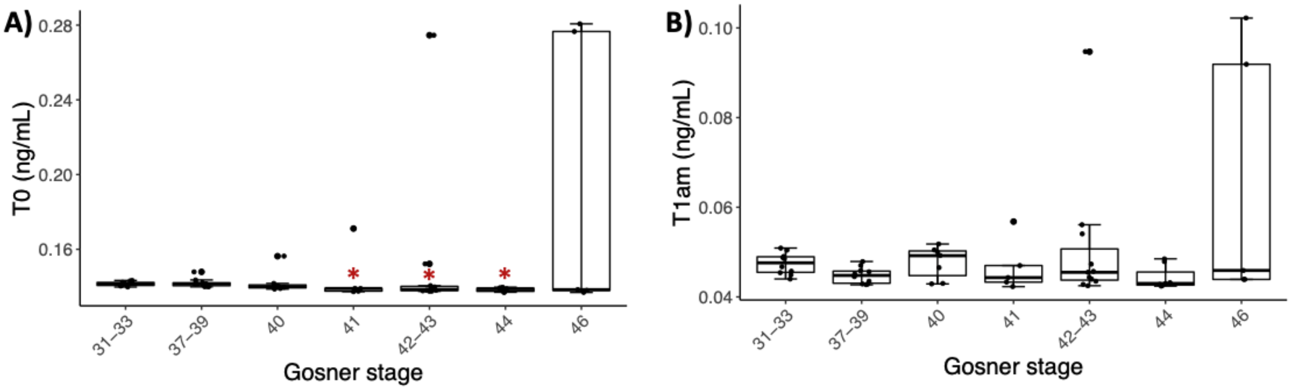


**Figure S1. Box plots showing serum concentration (ng/mL) of (A) T0 and (B) T1Am for the grouped Gosner stages.** Circles show individual replicates. Asterisks indicate significant difference from the premetamorphic Gosner stage 31-33 (Dunn’s post hoc test, p-value <0.05).

**Table S3. Spearman's rank correlations between gene abundance and TH concentrations**

| | ***thra*** | ***thrb*** | ***thibz*** |
|---|---|---|---|
| **T4** | rho = -0.09<br>p-value =0.49 | rho = 0.66<br>p-value =1.7e-08* | rho = 0.75<br>p-value =1.3e-11* |
| **T3** | rho = -0.26<br>p-value =0.04* | rho = 0.44<br>p-value =0.0003* | rho = 0.63<br>p-value =5.1e-08* |
| **T2** | rho = -0.2<br>p-value = 0.1 | rho = 0.32<br>p-value = 0.01* | rho = 0.32<br>p-value =0.01* |

### 3 Metabolomics quality assurance

A PCA plot including 8 composite quality control (QC) samples (Figure S2) shows that these are all centrally placed in the plot, which verifies the quality of the analysis sequence and instrument performance.

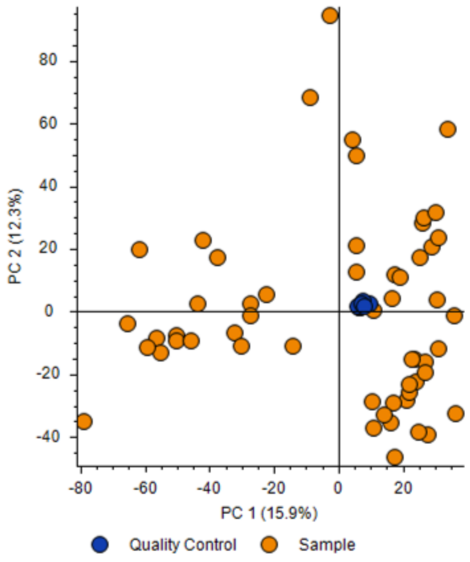


**Figure S2 Principal component analysis (PCA) of metabolomics data showing samples (yellow) and quality control samples.**

### 4 Multivariate analysis by sPLS-DA

Sparse partial least square discriminant analysis (sPLS-DA) was performed to discover biomarkers of the highest predictive power (Figure S3A). It is a linear, multivariate model which uses the PLS algorithm to allow classification of categorically-labelled data. sPLS-DA identifies components that best separate the sample groups and selects variables that best discriminate between groups. The first component explained 45% of the variation and metabolites separating prometamorphic and metamorphic climax (Gosner stage 44) animals. After tuning the model, 20 variables (i.e., metabolites) were selected on component 1. Figure S3B shows a plot of the loadings with a color assigned to each metabolite based on the sample group for which the mean area is maximum. This shows that all selected variables on component 1 have the highest abundance in Gosner stage 44 animals (teal, Figure S3). Table S4 shows the details for the 20 compounds including their stability (i.e., the proportion of cross validation folds (across repeats) upon which selection was based for a given component). The ID level is given as well as chemical class and name (tentative for all ID-level<1).

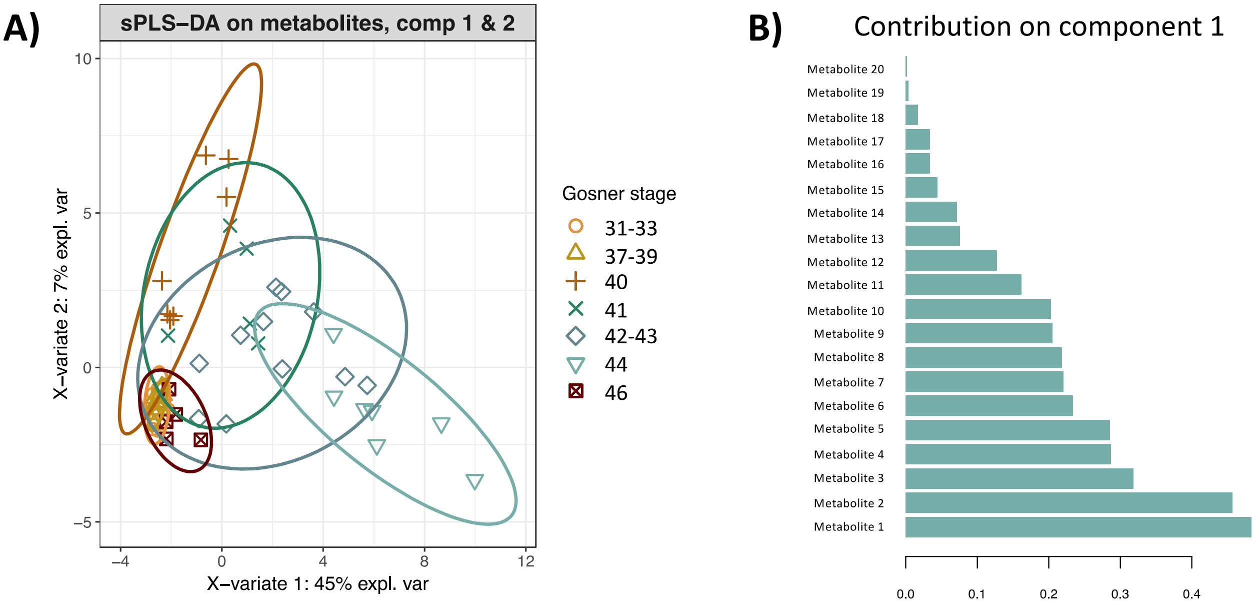


**Figure S3. (A) Sparse partial least square discriminant analysis (sPLS-DA) plot of the first two components with 95% confidence level ellipses. Samples are colored and grouped by Gosner stage. (B) The contribution of each metabolite selected on the first component (20 metabolites) with contribution ranked from bottom (important) to top. The color assigned to each metabolite is based on the sample group for which the mean area is maximum.**

**Table S4. Details for the 20 metabolites selected in component 1.** This includes their stability (i.e., the proportion of cross validation folds (across repeats) upon which selection was based for a given component). The chemical class, ID level and name, together with either a Metfrag score or a SIRIUS-4 formula score, and total explained intensity score are shown to demonstrate the level of confidence in the annotation. In addition, parameters from compound discoverer including the assigned formula, calculated mass, delta mass, mass-to-charge ratio (m/z), reference ion, and retention time (RT) are given.

| # | sPLS-DA stability score | Chemical class | ID level | Name | Metfrag score (2/2) | SIRIUS-4: formula score (%) | SIRIUS-4: Total explained intensity (%) | Formula | Calculated mass | Delta mass[3] (ppm) | m/z | Reference ion | RT |
|---|---|---|---|---|---|---|---|---|---|---|---|---|---|
| 1 | 1 | Long-chain fatty acid | 3 | | | 100 | 92.5 | C18 H28 O3 | 292.20362 | -0.77 | 293.2109 | [M+H]+1 | 20.609 |
| 2 | 1 | Linoleic acids and derivatives | 3 | 9-OxoOTrE | | 100 | 92.5 | C18 H28 O3 | 292.20359 | -0.87 | 293.21087 | [M+H]+1 | 20.274 |
| 3 | 0.77 | Long-chain fatty acid | 3 | | | 100 | 75.5 | C18 H28 O3 | 292.20363 | -0.72 | 293.21091 | [M+H]+1 | 20.491 |
| 4 | 0.9 | *Not possible to assign*[1] | 4 | | | | | C18 H28 O4 | 308.19854 | -0.72 | 309.20581 | [M+H]+1 | 20.998 |
| 5 | 0.8 | *Not possible to assign* | 4 | | | | | C22 H41 N O5 S2 | 463.24258 | -0.08 | 464.24985 | [M+H]+1 | 20.274 |
| 6 | 0.73 | *Not possible to assign* | 4 | | | | | C22 H41 N O5 S2 | 463.24261 | -0.02 | 464.24988 | [M+H]+1 | 20.609 |
| 8 | 0.83 | *Not possible to assign* | 4 | | | | | C28 H47 N3 O9 S2 | 633.27561 | 0.37 | 634.28288 | [M+H]+1 | 11.182 |
| 9 | 0.57 | Acyl carnitines | 3 | (5E)-tetradecenoyl-L-carnitine | | 99.1 | 93.1 | C21 H39 N O4 | 369.28766 | -0.69 | 370.29493 | [M+H]+1 | 16.491 |
| 10 | 0.83 | Acyl carnitines | 1 | Dodecenoylcarnitine | | | | C19 H35 N O4 | 341.25637 | -0.69 | 342.26365 | [M+H]+1 | 15.275 |
| 11 | 0.67 | Acyl carnitines | 3 | Undecylcarnitine | | 96.1 | 91.1 | C18 H35 N O4 | 329.25658 | -0.09 | 330.26385 | [M+H]+1 | 14.94 |
| 12 | 0.87 | Medium-chain hydroxy acids and derivatives | 3 | | 2 | | | C19 H32 O5 | 340.22481 | -0.49 | 341.23208 | [M+H]+1 | 17.26 |
| 12 | 0.47 | Medium-chain hydroxy acids and derivatives | 3 | | 2 | | | C19 H32 O5 | 340.22486 | -0.32 | 341.23214 | [M+H]+1 | 17.131 |
| 13 | 0.5 | *Not possible to assign* | 3 | | | | | C18 H28 O4 | 308.19856 | -0.72 | 309.20583 | [M+H]+1 | 18.494 |
| 14 | 0.43 | Linoleic acids and derivatives | 3 | 2R-HpOTrE | 1.99 | | | C18 H30 O5 | 326.20907 | -0.79 | 309.20578 | [M+H-H2O]+1 | 18.492 |
| 15 | 0.37 | Prostaglandin | 3 | Prostaglandin C1 | | 100 | 88.9 | C20 H32 O4 | 336.22975 | -0.93 | 337.23702 | [M+H]+1 | 19.77 |
| 16 | 0.37 | *Not possible to assign* | 4 | | | | | C22 H41 N O5 S2 | 463.2426 | -0.03 | 464.24988 | [M+H]+1 | 20.493 |
| 17 | 0.4 | Acyl carnitines | 1 | Decanoyl-L-carnitine | | | | C17 H33 N O4 | 315.24082 | -0.45 | 316.24809 | [M+H]+1 | 14.639 |
| 18 | 0.33 | Lactone | 3 | | 1.8 | | | C19 H30 O4 | 322.21419 | -0.68 | 323.22147 | [M+H]+1 | 18.312 |
| 19 | 0.37 | *Not possible to assign* | 4 | | | | | C21 H35 N O4 | 365.25637 | -0.65 | 366.26365 | [M+H]+1 | 15.271 |

| | | | | | | | | | | | | | |
|---|---|---|---|---|---|---|---|---|---|---|---|---|---|
| 20 | 0.87 | Cyclic alcohols and derivatives | 3 | | | 100 | 99.2 | C15 H22 O4 | 266.15168 | -0.47 | 267.15896 | [M+H]+1 | 18.331 |

[1]No $MS^2$

[2]Pattern coverage (%) displays the summed intensity of the matching isotope peaks in the measured MS1 spectrum relative to the summed intensity in the theoretical isotope pattern, thereby giving a quantitation of how well the measured isotope pattern matches the theoretical pattern.

[3]Delta mass (ppm) is the difference between the measured $M_w$ and the theoretical $M_w$ for the predicted composition
